# Improved antiproliferative activity of doxorubicin-loaded calcium phosphate nanoparticles against melanoma cells

**DOI:** 10.1101/2022.04.21.489118

**Authors:** Igor Barbosa Lima, Betania Mara Alvarenga, Priscila Izabel Santos de Tótaro, Fernanda Boratto, Elaine Amaral Leite, Pedro Pires Goulart Guimaraes

**Affiliations:** Federal University of Minas Gerais

**Keywords:** Senescence, drug delivery, cytotoxicity, A-375

## Abstract

The high incidence of melanoma has received significant attention. Despite advances in early detection and standard treatment options, new strategies that improve therapy with reduced side effects are highly desirable. Several studies have demonstrated the efficiency of doxorubicin (Dox) to treat melanoma, however, side effects limit its clinical use. Drug delivery systems, especially nanostructured ones, are a useful approach to enhance antitumor activity and reduce the toxicity of drugs. Here, we report the use of calcium phosphate nanoparticles functionalized with Dox and hyaluronic acid (N-Dox) to enhance Dox antiproliferative activity. The effects were accessed in A-375 melanoma cells, in which N-Dox significantly decreased IC_50_ over 48 hours (0.142 ± 0.07) compared to the free drug (0.44 ± 0.25). Treatment triggered DNA damage, increased nuclear area, and senescent phenotype. Furthermore, it did not form colonies after 14 days of incubation preceded by short exposure treatment. These preliminary results indicate that N-Dox hold promise for melanoma treatment, reducing the minimum effective dose and perhaps a reduction in the cost of treatment.

## 1. Introduction

Cancer remains a major issue to public health, being the second leading cause of death in the U.S., exceeded only by cardiac failure ^[1]^. Melanoma is a class of skin cancer that develops quickly, invasively ^[2]^ and, its occurrence is increasing over the years. The American Cancer Society informs that at least 106,110 new cases and 7,110 deaths from melanoma are expected to be reported in 2021^[3]^.

Most melanoma cases are associated with a mutation that results in overexpression of B-RAF proto-oncogene, concomitant with a reduction of p53 activity ^[4]^. BRAFq enzyme acts as a modulator for cell cycle progression under external stimulus (e.g., growth factors and hormones). However, once the BRAFq gene expression is upregulated, this cell division pathway is triggered without external stimuli, which leads to uncontrolled cell proliferation. Due to this, the BRAFq pathway has become a target for melanoma treatment, focusing on the development of drugs that act directly by blocking its activity ^[5]^.

On the other hand, some melanoma cases do not present BRAFq upregulation, which brings a research approach to alternative therapeutic targets ^[6]^. Doxorubicin hydrochloride (Dox) is not currently used as a treatment protocol for melanoma ^[7]^, however, several studies demonstrated Dox’s efficacy to treat it ^[8–10]^. Despite the advances, clinical use of Dox is still a challenge because of a range of serious side effects occurring during treatment (nausea, headache, etc.) and long-term sequelae ^[11]^ that include marked dose-dependent cardiotoxicity and nephrotoxicity ^[12,13]^.

In this context, the reduction of the effective dose becomes a therapeutic target of interest that can be achieved by nanotechnology ^[14]^. Calcium phosphate nanoparticles (NPC), are composed of magnesium, calcium, and phosphorus, have amorphous nature, spherical shape, an average diameter of 149 nm, and are efficient in delivering drugs to different types of cells, such as macrophages and breast cancer cells ^[15,16]^.

NPC can be functionalized by hyaluronic acid (HA) as a strategy to increase targeting to cancer cells ^[17]^. HA binds with the CD44 receptor ^[18]^, which is overexpressed in several cancers, including melanoma ^[19,20]^. Besides that, NPC cad adsorb Dox (approximately 17.72%), which decreases zeta potential ^[16]^. These characteristics are interesting to increase tumor retention by enhanced permeability and retention effect ^[21]^. These findings are consistent with the antiproliferative activity of Dox-loaded NPC observed against human breast cancer cells ^[16]^. However, the possibility of being used as a therapeutic platform for the treatment of melanoma has not yet been investigated. Based on that, we hypothesize that using Dox-loaded NPC for drug delivery potentiates doxorubicin activity against melanoma cells.

We performed Dox adsorption on NPC followed by functionalization with hyaluronic acid, resulting in a formulation mentioned as N-Dox. N-Dox cytotoxic activity against melanoma cells (A-375 cell line -Homo sapiens) was accessed, as well as the investigation of the cytotoxicity mechanism.

## 2. Material and methods

### 2.1. Synthesis of NPC and N-Dox preparation

NPC was synthesized using phosphate salts solutions at controlled Ph in a semipermeable membrane system, 25 mm wide (MWCO 15,000 Spectrum Medical Industries, Inc., Fisher Scientific, USA) as described by Alvarenga^15^. A suspension containing 0,4 mg of NPC was eluted to 380 µL of Milli-Q water, mixed to 20 µl of a 1.8 mM Dox (Sigma Aldrich, USA), and homogenized on vortex for 5 minutes to lead to Dox adsorption on NPC surface by self-assembly, purified through three consecutive cycles of washing and centrifugation^[22]^, and functionalized by 200 µl of a 3% (v/v) solution of hyaluronic acid (HA) (Merck, Germany) at room temperature. NPC samples were functionalized only with HA and used as a control ^[16]^.

### 2.2. Dox loading and adsorption efficiency

The amount of Dox loaded on N-Dox was determined by high-performance liquid chromatography (HPLC) with a fluorescence detector as described by Boratto^[23]^. Three independent preparations were analyzed. To measure the concentration of Dox adsorbed, the pellet was resuspended in 200 µl of HCl at 0.5 mM, since acidification digests the material. The analyses were performed in a Waters chromatographer (Waters Instruments, 1200 series, Milford, USA). The percentage of the load was calculated based on the equation Eq. (1):

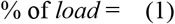

where *c* is the concentration of Dox in the sample in µM and 90 is the initial concentration in µM.

### 2.3. *In vitro* drug release studies

To determine the release rate of doxorubicin from N-Dox, we use the dialysis bag technique as described by Krai ^[24]^. 400 µl of N-Dox suspension was prepared in triplicate and after 0.5; 1; 3; 4; 6; 8; 20 hours, samples with 200 µl were removed and the same volume of PBS was replaced to avoid an increase in the concentration of the medium. The aliquots were analyzed by HPLC as previously described.

### 2.4. Cell culture

All cell lines were cultivated in DMEM medium (Sigma Aldrich, USA), containing 10% fetal bovine serum (GIBCO BRL, Grand Island, NY, USA) and 1% antibiotic solution100 IU. mL^-1^ of penicillin and 100 µg. mL^-1^ of streptomycin (GIBCO BRL, Grand Island, NY, USA) and kept in a humidified atmosphere with 5% CO_2_ at 37 ° C. A375 and HEK-293 cell lines were kindly donated by Dr. Helen Lima del Puerto/UFMG – Brazil. Cells were authenticated and tested for infection by *Mycoplasma sp*. through PCR.

### 2.5. Cytotoxicity assessment

Cell proliferation was assessed by the colorimetric assay of sulforhodamine B 2-(3-diethylamino-6-diethylazaniumylidene-xanthen-9-yl)-5-sulfo-benzenesulfonate (SRB) (Sigma Aldrich, USA)^[25]^. A-375 cells were seeded at a density of 5×10^3^ cells per well in 96-well plates and treated with NPC, N-Dox, and free Dox by using eight serial dilutions (1: 5) with concentrations between 120 - 0.0 µg. mL^-1^, 1.42 - 0.0 µM, and 180 - 0.0 µM. respectively. The spectrophotometric reading of the OD was performed at 510 nm in a multiwell plate reader (Varioskan Lux - Thermo Fisher Scientific^®^, MA, USA).

Results were expressed as a percentage of cell viability as a function of concentration. The OD was normalized as the percentage of cell viability (% CV) considering the absorbance of the PBS-treated control (OD_PBS_) as 100% viability, according to the equation Eq. (2):

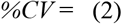

Values of inhibitory concentration for 50% of the cell population (IC_50_) were calculated by non-linear regression using GraphPad Prism^®^ software version 5.0 (GraphPad Software Inc USA). The selectivity indices (SI) were calculated for each formulation as the ratio between the IC_50_ of the HEK293 cell line and the IC_50_ of the A375 cell line _[26]_.

### 2.6. DNA quantification and evaluation of cell cycle

The quantification of DNA and analysis of the cell cycle were performed as described by Riccardi and Nicoletti ^[27]^. Subsequently, cells were treated with concentrations referring to the 48 hours IC_50_ (Dox, N-Dox, and NPC at 0.45 µM, 0,15 µM, and 0,6 µg. mL^-1^ respectively). After 48 hours of treatment, cells were stained with hypotonic fluorochromic solution (HFS) and parameters were acquired by flow cytometry (FACSCan^®^, BD Biosciences, USA) and analyzed in the software FlowJoX.0.7^®^ (Tree Star, Inc.USA).

### 2.7. Clonogenic assay

The long-term survival of A375 cells after treatment with the formulations was accessed through the clonogenic assay ^[28]^. The cells were treated as described in topic 2.6., Briefly, plates were incubated for 14 days and then fixed with 4% p-formaldehyde, washed, and stained with 0.1% violet crystal. Only colonies with at least 50 cells were considered and counted manually. The results are expressed as a fraction of survival, obtained by the equations below. Plating efficiency (PE) was determined as the ratio between the number of colonies formed in the well without the treatment and the number of cells seeded, multiplied by 100 Eq. (3). The surviving fraction (SF) was calculated by dividing the number of colonies formed after treatment by the number of seeded cells multiplied by PE Eq. (4).

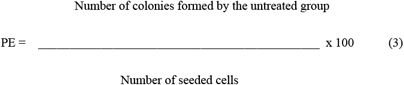

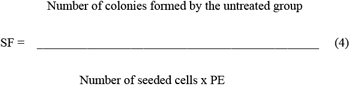

### 2.8. Nuclear morphometric analysis (NMA)

A375 cells were seeded at a density of 5×10^4^ cells per well in 8 well Tek Chamber Slides (Nunc™, Thermo Fisher, USA) and incubated for 24 hours for stabilization. Cells were treated as described in topic 2.8. and stained with Hoechst reagent (NucBlue™ Live ReadyProbes™, USA) according to the manufacturer’s instructions. The evaluation of nuclear morphometry was carried out by measurement of nuclear shape and area. These variables were used to calculate the nuclear irregularity index (NII), which is used to indicate the mechanism of cell death induced by treatments. The images were obtained in a confocal microscope (LSM 880 ZEISS, Germany) (magnification x 400) and analyzed using ImageJ software as described by Filippi-Chiella^[29]^.

### 2.9. Statistical analysis

All experiments were carried out at least in triplicate. Data were expressed as mean ± S.D. (standard deviation). Statistical analyzes were performed using GraphPad Prism software, version 5.0, GraphPad Software, Inc, USA. The normality of the data was ascertained by the Kruskal-Wallis test and the difference between means of treatments was accessed by One-way ANOVA with Tukey’s multiple comparison test. Significant differences were considered at the level of p ≤ 0.05.

## 3. Results and discussion

### 3.1. Characterization of nanoparticles

NPC used in this work was loaded with Dox through electrostatic interactions, in an aqueous medium to promote self-assembly and previously characterized ^[16]^. It is a process less laborious, less costly, and avoids the use of organic solvents (such as dimethyl sulfoxide) that can increase the toxicity of formulations^30^. The coating of the nanoparticles with HA also was carried out by adsorption, and this technique has been described as satisfactory for this purpose ^[31]^.

Dox quantification by HPLC was performed to confirm the loading observed previously. N-Dox showed a mean Dox concentration of 14.2 ± 4,8 µM (this value was considered as the average concentration of Dox present in N-Dox for the subsequent experiments and calculations), regarding a mean load of 15.78 ± 5.33% (figure 1. a), consistent with the values found previously^[16]^. Positively charged cerium oxide nanoparticles, with zeta potential ranging from -31.16 to 40.2 mV, can load to bovine serum albumin (negatively charged protein) since the zeta potential is positive^[32]^. The loading of Dox on the NPC in our study follows a similar pattern: negatively charged NPC (negative zeta potential) performs electrostatic interactions with Dox, a cationic compound^[33]^.

**Figure 1.**
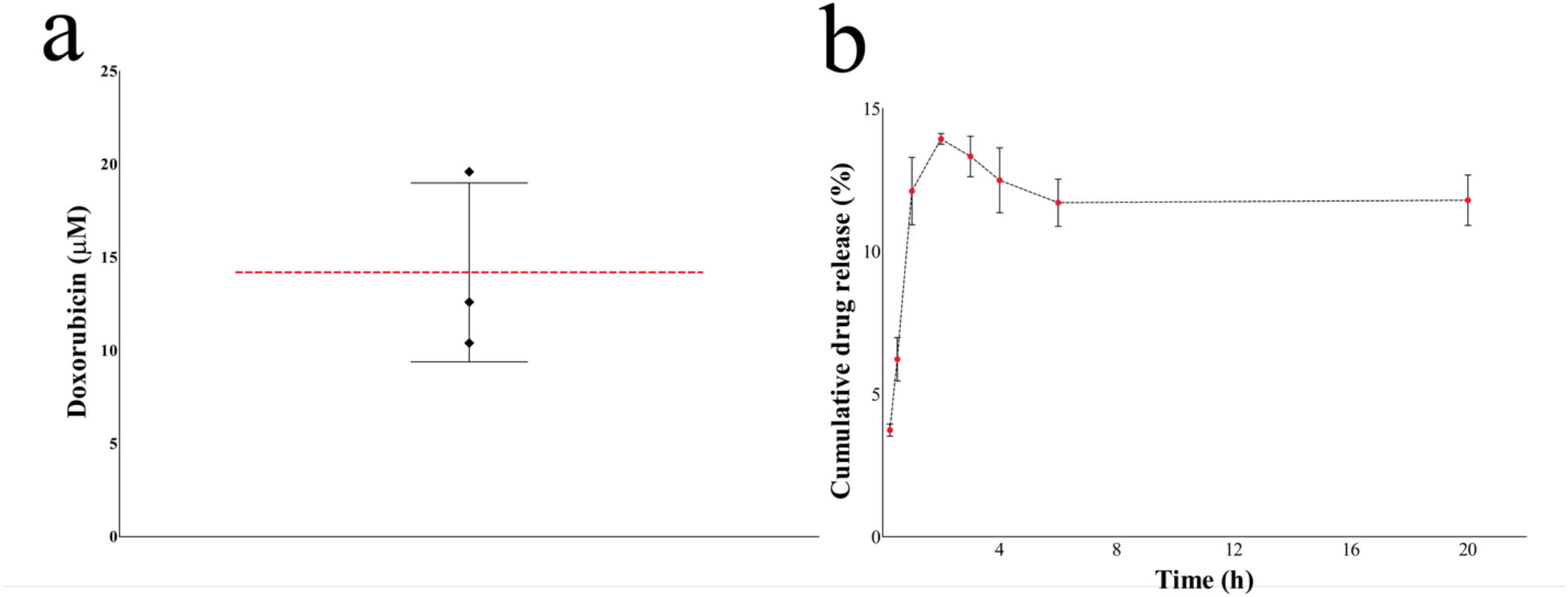
The concentration of Dox adsorbed on NPC. Analysis performed on HPLC after acid digestion (a); release/diffusion profile from N-Dox. Initial concentration 25,2 mg. mL^-1^ (b). Red dashed lines represent the average of the measurements, n = 3, ±S.D.

The diffusion/release profile of Dox from N-Dox (figure 1. b) shows a burst of release between 1 and 2 hours of incubation at pH 7.4, with a peak in approximately 2 hours (13.9 % ± 0.18). This finding suggests that N-Dox is a drug delivery system with a slow-release rate, which likely provides constant drug release in the first hours, increasing the drug retention time. Nano formulations are expected to increase the drug retention time of Dox^[34]^ and can influence the enhanced permeability and retention effect^[35]^.

### 3.2. Cytotoxic activity

Dox can act in both the cytoplasm and nucleus^[36]^. After its internalization, Dox is reduced to an unstable semiquinone that is transformed again into Dox, generating reactive oxygen species (ROS)^[37]^. On the other hand, Dox reach the nucleus, causes damage to topoisomerase II, and alters the expression of genes involved in DNA repair and cell cycle regulation^[38]^.

A-375 cells presented remarkable morphological changes after treatments (figure 2.a), consistent with dose-response cytotoxicity curves (figure 2.b). Treatment with N-Dox for 48 hours significantly reduced the dose required (0.142 ± 0.07) compared to the free drug (0.44 ± 0.25). After 48 hours the cytotoxic effects of N-Dox increase, once this stage, there is a damage accumulation caused by Dox molecules slowly released from the nanoparticles, causing DNA damage (figure 3. a). It was not possible to calculate the IC_50_ of unloaded NPC since there was no significant reduction in cell viability after treatment.

**Figure 2.**
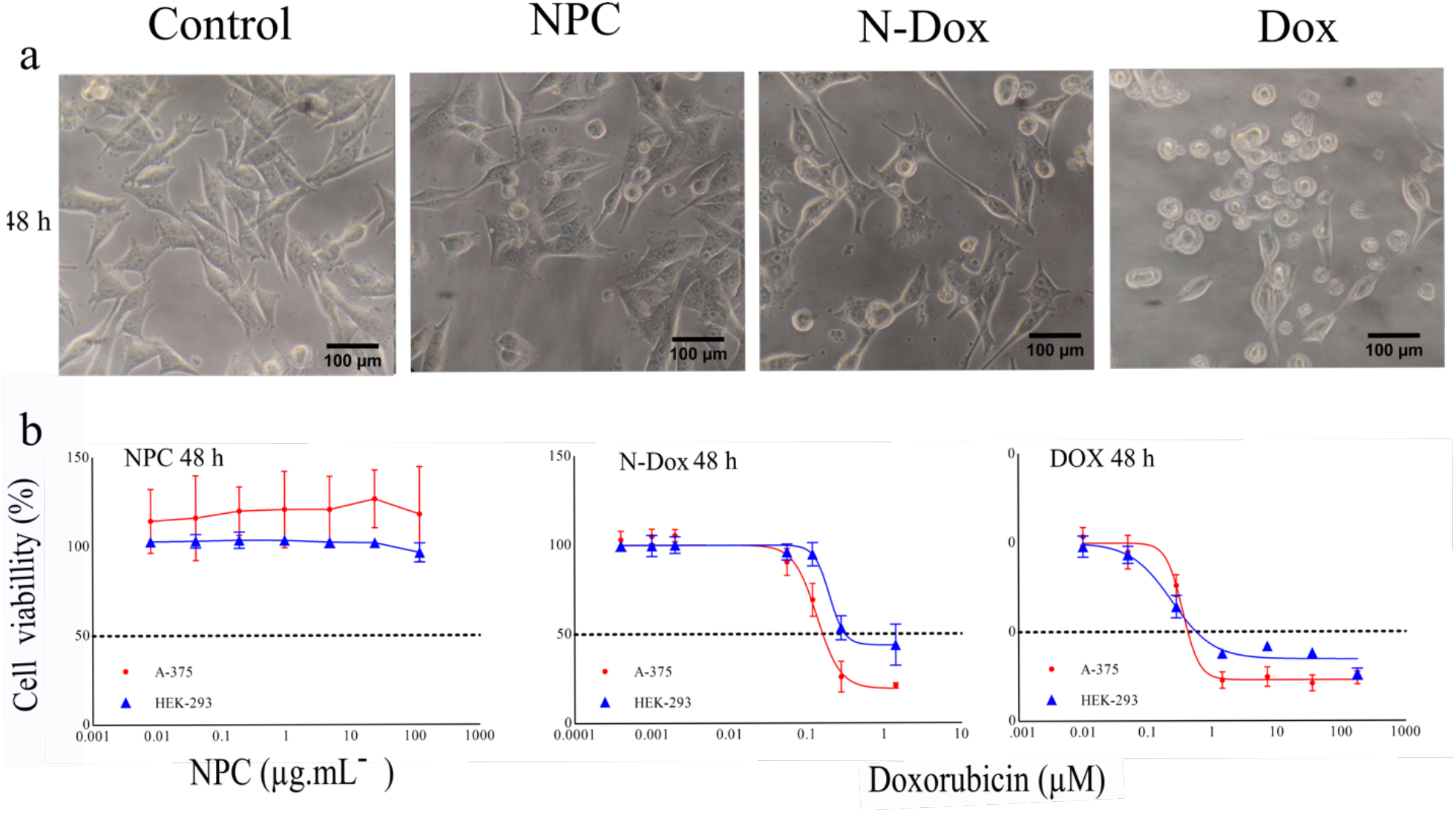
Antiproliferative activity of NPC, N-dox, and Dox after 48 hours. Representative photomicrographs of the morphology of A-375 cells after treatment obtained under a phase-contrast microscope, scale bars: 100 µm (a); Dose-response curves of cell lines A-375 and HEK-293 after treatment with Dox, N-Dox, and NPC for 24, 48 e 72 hours (b).

**Figure 3.**
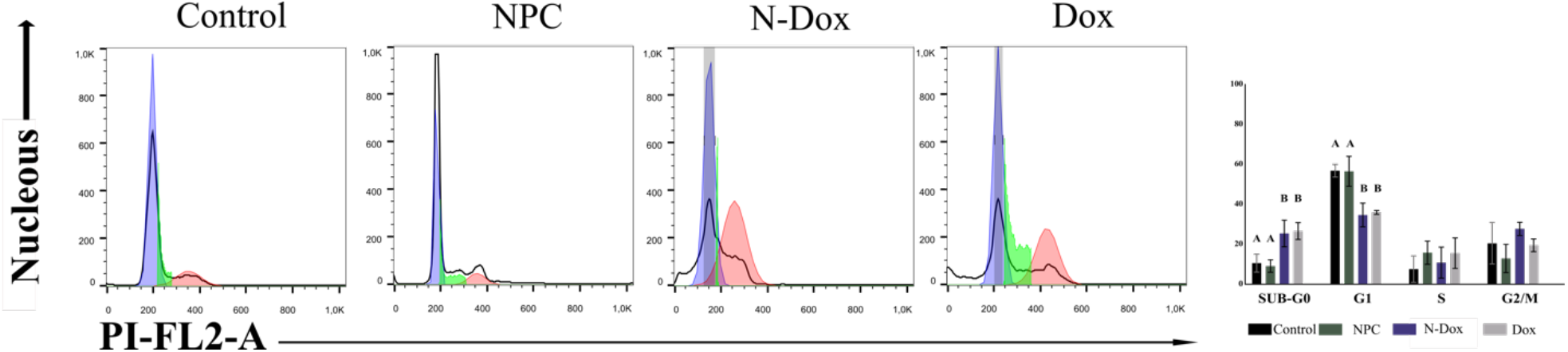
Effect of treatment with Dox (0.45 µM), NPC-Dox^+^ (0.13 µM); and NPC (0.6 mg.ml-1) to cell cycle of A-375 cells. Data expressed as mean ± SD obtained from three independent experiments carried out in duplicate, n = 6. Data were compared by ANOVA and Dunnett’s means test with a significance level at p <0.05. The letters A and B show a significant difference between treatments.

*In vitro* models are efficient to access the security of new technologies and drive research promoting the policy of the three R’s (replacement, reduction, and refinement) for using animals in research^[39]^. HEK293 cells are a classic model for predicting drug safety and nephrotoxicity^[40]^. The selectivity indices (SI) (Dox = 0.54; N-Dox = 3.2) shows that N-Dox have selective cytotoxicity. Since health cells are less affected by N-Dox, that could represent reduced side effects hereafter, when it is *in vivo testing*.

### 3.3. Cell cycle assessment

To determine the predominant cell death mechanism over melanoma cells, the viability must be evaluated in line with cell cycle progression. Treatment with NPC did not cause any change in the cell cycle. Treatment with N-Dox (0,15 µM) and Dox (0,45 µM) in A375 cells induces significant changes in cell cycle progression after 48 hours of treatment (figure 3. b). We observed a significant increase in DNA damage which is a strong indication of the occurrence of programmed cell death^[41]^ and a reduction in the number of diploid cells (G0/G1). There was no alteration in the number of cells in the S phase. Since no significant difference in cell cycle progression was observed after treatment with N-Dox, compared to the drug-free form, it is presumable that loading doxorubicin to NPC as a platform for delivery, does not alter the Dox mechanism of action.

### 3.4. Cell death pathway

The occurrence of tumor recurrence and multiple drug resistance is characterized by resistance to several drugs with different treatment targets^28^. The clonogenic assay simulates the effect of long-term treatment and tests the ability of a tumor cell to perform infinite cell divisions, even after exposure to a cytotoxic stimulus^[28]^. Our results showed that N-Dox administration was able to prevent the formation of A375 cell colonies after 14 days of incubation, which indicates that the cells were not able to reverse the damage caused by the short exposure period treatment^28^. NPC treatment formed colonies as well as control. Plating efficiency (PE) and surviving fractions are summarized in table 1.

**Table 1.**
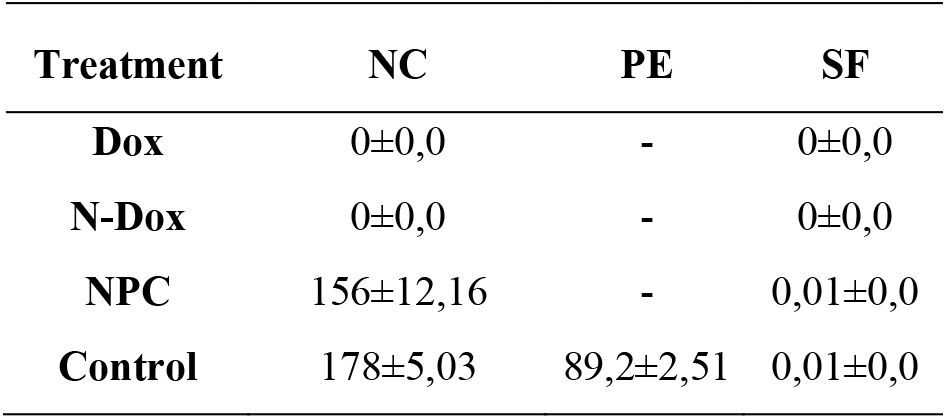
Number of colonies (NC), plating efficiency (PE), and a surviving fraction (SF) of A375 cells after treatment. Data expressed as mean ± S.D. n = 3.

| **Treatment** | **NC** | **PE** | **SF** |
| --- | --- | --- | --- |
| **Dox** | 0±0,0 | - | 0±0,0 |
| **N-Dox** | 0±0,0 | - | 0±0,0 |
| **NPC** | 156±12,16 | - | 0,01±0,0 |
| **Control** | 178±5,03 | 89,2±2,51 | 0,01±0,0 |

To ascertain which cell death mechanism is activated by N-Dox, cell nuclei were stained by Hoechst (figure 4. a) for subsequent Nuclear Morphometric Analysis^[29]^. Both Dox and N-Dox triggered the occurrence of large and regular nuclei in A375 cells (64.2% ± 2.7 for Dox and 68.8% ± 12.8 for N-Dox) i.e., senescent cells (figure 4. b). Cancer cells can enter a state of senescence after being exposed to chemotherapeutic agents^[17,42]^ e.g., doxorubicin, which is a potent inducer of senescence in cancer cells^[43]^. This phenomenon is a desirable form of treatment for cancer known as therapy-induced senescence (TIS) as it stops the progression of tumors less aggressively^[44]^, which is considered a good strategy to stop melanoma development^[45]^. The ability to form colonies was completely inhibited suggesting the absence of cells capable of evading the senescence mechanism^[46]^ since “scaper” cells are more invasive and form colonies faster^[47]^.

**Figure 4.**
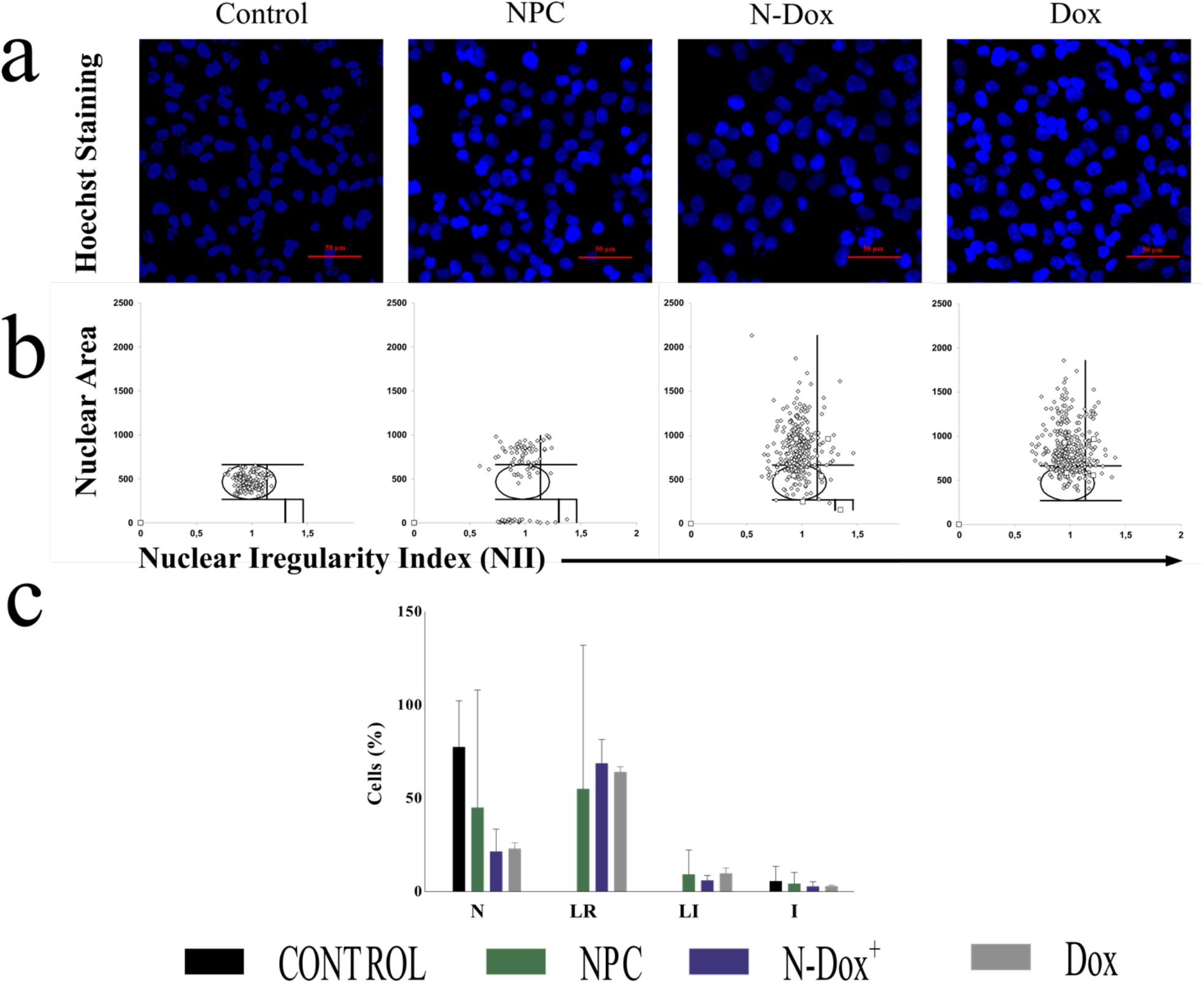
Nuclear Morphometric Analysis of A-375 cells treated as indicated. (a) Nucleus was stained with Hoechst for confocal microscopy imaging; scale bars, 50 µm; magnification, 400 x, and have their nuclear area (y-axis) and shape (x-axis) measured.). (b)Nuclei classified as N = normal; I = Irregular; LR = Large Regular, and LI = Large Irregular, showing the percentage of senescent nuclei (LR). Results were compared by ANOVA and Tukey’s mean comparison with a significance level at *p* <0.05. Letters A and B indicate a significant difference between treatments, n = 6.

## 4. Conclusion

NPC has shown to be an efficient drug delivery system for Dox with a mean retention capacity of 15.78 % ± 5,33 and, has a slow-release rate, which could provide a longer drug retention time in the circulation. N-Dox provided potentiation of Dox effect after 48 hours of treatment and induced senescent phenotype. As no colonies were formed after the long-term survival assay, we suggest the absence of “scaper” cells. Our preliminary findings demonstrated that N-Dox has enhanced activity compared to Dox. The quantification of the expression of the genes involved with the progression of the cell cycle is fundamental to confirm the absence of escapers cells. This approach will allow that the formulation has its effectiveness and safety assured. N-dox is promising as a potential treatment for melanoma that could reduce damage to patients and reduce treatment costs due to the smaller amount of drug used.

I would like to thank everyone who directly or indirectly contributed to this research.

## Author contributions

Igor Barbosa Lima – execution of all experiments, analysis of results, and writing of the manuscript.

Betania Mara Alvarenga – synthesized the nanoparticles and helped in carrying out the experiments.

Priscila Totaro – patent holder and postulated the methodology and manuscript review.

Fernanda Boratto - participated in the planning and execution of HPLC analyses.

Elaine Amaral Leite - participated in the planning and execution of HPLC analyses. Pedro Pires Goulart Guimaraes - participated in the planning and execution of the nuclear morphology analysis.

## Ethics approval

This is an observational study. The XYZ Research Ethics Committee has confirmed that no ethical approval is required.

